# Lipid Demixing Reduces Energy Barriers for High Curvature Vesicle Budding

**DOI:** 10.1101/2024.10.24.620077

**Authors:** Itay Schachter

## Abstract

Budding relies on membrane asymmetry, including composition, area, and osmotic differences, and involves large curvature changes in nanoscale lipid vesicles. So far, the combined impact of asymmetry and high curvatures on budding has remained unknown. Here, using continuum elastic theory, the budding pathway is detailed under realistic conditions. It shows that budding is less favored in smaller vesicles but lipid demixing can significantly reduce its energy barrier and yet high compositional deviations of more than 7% between the bud and vesicle only occur with phase separation on the bud.

---

Nanoscale budding plays a crucial role in key biological processes such as endocytosis, viral budding [1] and intraluminal vesicle formation [2]. Budding is defined as a process wherein a single closed membrane undergoes a sequence of shape changes, ultimately leading to the formation of a distinct daughter vesicle (DV) connected to the original membrane via a narrow neck (Fig. 1A). Here, the closed membrane is set as a spherical or prolate mother vesicle (MV). Budding may become energetically favorable whenever some asymmetry arises between the outer and inner leaflets (Fig. 1B), such as in lipid or solution composition, area, osmotic conditions or a combination of these [3]. These asymmetries are prevalent in cells and organelles [4–7], with transmembrane proteins such as lipid transporters and channels, modulating the membrane asymmetry and influence highly-regulated biological processes involving budding [8,9]. Thus, understanding the impact of asymmetry on budding may enhance our comprehension of the regulation of processes—such as viral budding, their pathologies and their sensitivity to asymmetry in general. The area-difference elasticity (ADE) model [10] specifically predicts that a combination of area asymmetry, lower than ideal vesicle water content and a positive bilayer spontaneous curvature— due to lipid or solution composition asymmetry—may induce outward budding. However, the combined contribution of the possible asymmetries to budding, especially at the nanoscale, has thus far remained enigmatic.

**FIG. 1.**
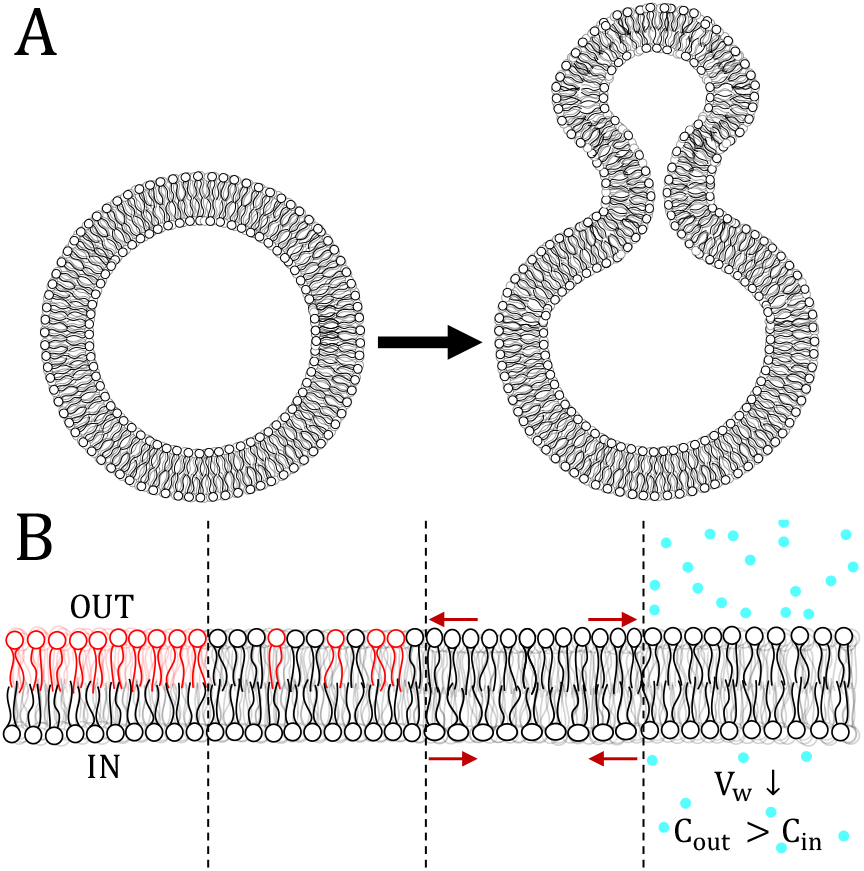
(A) Illustration, not to scale, of the initial (left) and final (right) states of outward budding. Initially, no significant curvature is present. In the final, budded state, high curvature appears at the neck region and the DV. (B) Types of membrane asymmetries: 1. Composition: Completely asymmetric (left) or partially asymmetric (middle left), as seen in vesicles from layer-by-layer assembly and some outer-leaflet modification methods, respectively. 2. Leaflet area: Non-ideal lipid numbers in each leaflet cause opposite stresses (middle right). 3. Osmotic conditions: Hypotonic conditions reduce the volume of entrapped water (right).

For experiments, layer-by-layer assembly and outer-leaflet modification methods [11, 12], such as lipid exchange and enzymatic modification, enable the creation of asymmetrical model vesicles in terms of composition and possibly leaflet area. These methods allow the study of how asymmetry affects membrane properties and processes [13–17]. However, significant variations between vesicles within a sample limit the experimental accuracy. These variations are in the degree of composition asymmetry [11], vesicle size [18], and water content [19]. Also, imaging nanoscale budding requires techniques such as electron microscopy, which lack the temporal resolution to fully capture this dynamic process [20]. While molecular dynamics (MD) simulations offer the necessary spatial and temporal resolutions and allow for high control over parameters like the degree of asymmetry, they are currently restricted by their limited scales [21]. Even with these limitations overcome, relying solely on experiments and simulations without theoretical frameworks is insufficient for comprehending the forces driving budding. Thus, a theoretical description is required to fully understand this process.

In nanoscale vesicles, budding generates relatively high curvatures at the nanometer-sized DV as well as the neck region. The ADE model struggles in this scenario because it relies on the assumption that the membrane thickness is negligible compared to the vesicle radius. Furthermore, as was suggested in a recent experimental study [17], the high curvature differences may induce local lipid sorting, yielding substantially different compositions between the MV and DV. Membrane heterogeneity is unaccounted for by the ADE model and, to this author’s knowledge, has not yet been quantitatively described in the literature in the context of budding.

Here, I use a highly-detailed elastic model to investigate the budding process, which alleviates the limitations of previous experimental and theoretical studies. It describes a single nanoscale vesicle under relevant experimental and physiological conditions. The findings reveal that budding is more prohibited in these vesicles. Also, lipid heterogeneity, similar to that in cells and organelles, can lower the budding energy barrier, though significant lipid sorting requires domain formation.

Building on the formalism that extends the Helfrich– Hamm–Kozlov model [22–24], as detailed by Ryham et al. [25–27], this study presents a continuum model for an enclosed vesicle (Fig. 2A); Section S1 details the implementation. For each leaflet (*j* = *o, i*, denoting outer and inner), the lipid vector field 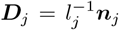 describes the local (relaxed) lipid length *l*_*j*_ (*l*_0_) and the unit lipid director normal ***n***_*j*_. This field, defined over the leaflet neutral surface *S*_*j*_, points from the neutral surface to the midplane surface *S*_*m*_. Cylindrical symmetry is assumed for these surfaces. The elastic contribution to the free energy for each leaflet, based on the modes of deformation shown in Fig. 2B, is given by

**FIG. 2.**
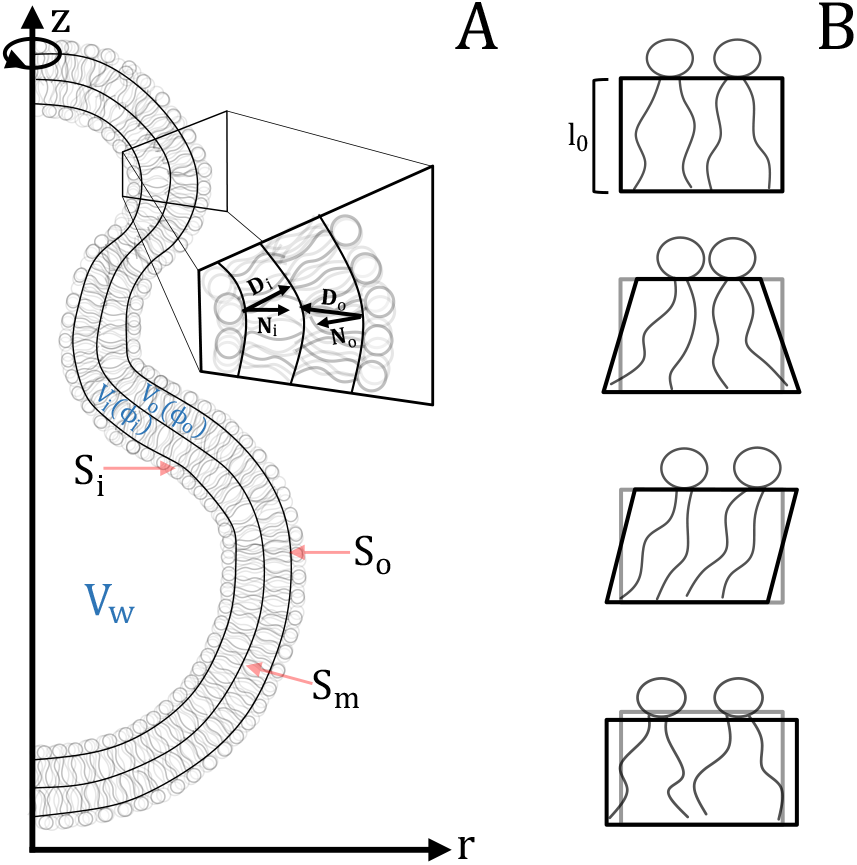
(A) Schematic view of a budded vesicle showing the vector fields described in the main text. (B) Schematic of the modes of deformation considered by the elastic theory, from top to bottom: undeformed, bend, tilt and area stretch.

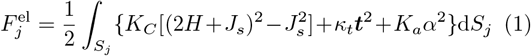

The first term represents the bending energy, with the bending modulus *K*_*C*_ determining the energy penalty for splay deformations, 2*H* =∇ · ***n***, from its natural value, defined as minus the spontaneous curvature *J*_*s*_ (positive for a spherical micelle). The second term describes the tilt energy governed by the tilt modulus *κ*_*t*_ and the tilt deformation ***t*** = ***n****/*(***N*** · ***n***) − ***N***, where ***N*** is the surface normal. The final term defines the area-compressibility energy with modulus *K*_*a*_ for deviations in the local areaper-lipid (APL), *a*, from its relaxed value, *a*_0_, described by 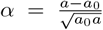. Assuming local lipid incompressibility (*al* = *a*_*0*_ *l* _*0*_), 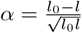.

The model includes volume constraints to accurately describe the physical and experimental conditions. Each leaflet’s volume is kept constant, reflecting global lipid incompressibility and the negligible rate of phospholipid flip-flop [28]. The volume of the entrapped water is also set constant, as it remains practically unchanged under specific osmotic conditions [3] and is influenced by them and by the vesicle fabrication method [19]. The model was validated under various scenarios (Section S1 A) and is available on GitHub.

Consider an initially spherical vesicle with a midplane radius, *R*_*m*_ = 50 nm. The difference in the (relaxed) area between the outer and inner leaflets is Δ*A* = 4*l*_0_*R*_*m*_*m*_0_, where *m*_0_ is a unitless factor accounting for leaflet area asymmetry. For no asymmetry, *m*_0_ = 4*π*; *m*_0_ is larger (smaller) than 4*π* when the outer leaflet has a lipid excess (deficiency) and vice versa for the inner leaflet. The entrapped water volume, *V*_*w*_, may deviate from its ideal value of 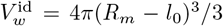, such that 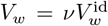, with *𝒱* defined as the relative water content factor. Using these two factors, *m*_0_ and *𝒱*, the equilibrium phase was determined, yielding a phase diagram (Fig. 3).

**FIG. 3.**
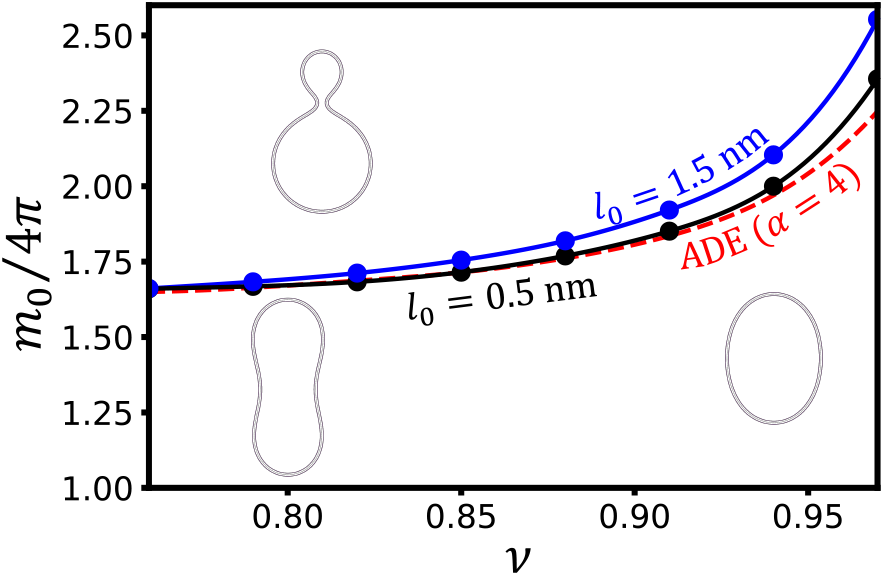
Phase diagram of the elastic model in the (*𝒱, m*_0_) plane involving budding. The coexistence curve between the prolate and budded states, interpolated with a cubic spline (solid lines) between sampled values (circles), shifts upward with monolayer thickness, *l*_0_. Other elastic parameters are *K*_*C*_ = 10 *k*_B_T, *κ*_*t*_ = 10 *k*_B_T nm^*−*2^, and 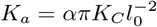 with *α* = 4. As *l*_0_ *→* 0, the curve should nearly coincide with the ADE model’s prediction from Ref. 10 for *α* = 4 (dashed red line); the curve shown was originally suggested for *α* = 1 (see Section S2).

The non-negligible monolayer thickness relative to the vesicle radius (*l*_0_ ≪ *R*_*m*_) makes budding less favorable because the coexistence curve in the (*ν, m*_0_) plane shifts upwards when a typical *l*_0_ of 1.5 nm is chosen. This shift is particularly pronounced for vesicles with nearly ideal water content volume (*ν* ≈ 1), making budding more inhibited in them. When *l*_0_ ≪ *R*_*m*_, the bending free energy density is described as a function of the midplane mean curvature, *H*, via *K*_*C*_(2*H*)^2^. For nanoscale vesicles, however, positive higher-order terms, linear in *l*_0_ [29], contribute enough to the bending free energy to somewhat inhibit budding.

The coexistence curve for the thinner membrane (*l*_0_ = nm) closely aligns with the one predicted by the ADE model. A comparison of the current model results with those provided in the original paper of the ADE model [10] reveals a minor but consistent miscalculation in that paper, further detailed in Section S2. Nevertheless, the ADE model provides a good approximation when *l*_0_*/R*_*m*_≈ 0.01 or lower, i.e., for vesicles with a radius of several hundred nanometers.

Here, the model is extended to include lateral compositional variation. For simplicity, we will focus on binary lipid mixtures. To this aim, a scalar field *ϕ*, representing the molar ratio of the additional lipid, is defined over *S*_*j*_. Consequently, the model now includes a mixing free energy contribution in mixed leaflets, relative to homogeneous mixing with *ϕ*_0_ = ⟨*ϕ*⟩. This contribution is expressed as 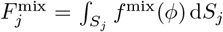, where *f* ^mix^(*ϕ*) is the mixing free energy density. Specifically, *f* ^mix^ is chosen as

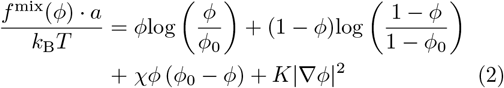

The first two terms describe the ideal mixing entropy between the components, while the third term introduces a non-ideal interaction between the two lipids, whose magnitude is dictated by an interaction parameter, *χ*. The last term details the penalty for spatial variations in the composition and ensures that *ϕ* remains continuous; further details in Section S1. In a planar membrane, phase separation can occur when *χ >* 2, because *f* ^mix^ then has two points of common tangent. Analysis of MD simulations shows that eq 2 effectively captures the extent of mixing for different binary mixtures over a wide range of *ϕ* values (Section S4). Similarly to the volume constraints, the average composition of mixed leaflets is maintained. Elastic deformations promote demixing whenever lipids have different physical properties [30–32]. The model only includes variations in *J*_*s*_, which is intimately related to lipid geometry [33], as the membrane shape is very sensitive to it [34]. It varies linearly between its values for the two components with the local composition; i.e., 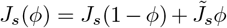, where 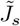 is its value for the additional lipid.

The default lipid parameters in the model for the subsequent analyses match those of 1-palmitoyl-2-oleoyl-*sn*-glycero-3-phosphocholine (POPC), and are specified as follows [32, 35]: *K*_*C*_ = 14.49 *k*_B_*T, κ*_*t*_ = 9.5 *k*_B_*T* nm^*−*2^, *K*_*a*_ = 28.93 *k*_B_*T* nm^*−*2^, *a*_0_ = 0.639 nm^2^, and *l*_0_ = 1.51 nm. The spontaneous curvature of POPC is negligible [36], and therefore *J*_*s*_ is set to zero.

The effect of lipid mixing on the budding free energy pathway is studied here, following experimental conditions involving outer leaflet lipid insertion [11, 17]. In this setup, 10% extra lipid volume is added to the outer leaflet of an initially unstressed, symmetric POPC vesicle with a radius of 50 nm and 96% relative water content volume, as vesicles usually contain less water than ideal [3, 19]. Furthermore, the outer leaflet is set as an ideal binary mixture (*χ* = 0) with an average spontaneous curvature 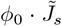 of 0.04 nm^*−*1^, which arises due to the presence of the additional lipid. The minimal free energy path for the budding of this vesicle is evaluated using the Nudged Elastic Band (NEB) method (details in Section S5) for different values of *ϕ*_0_ (Fig. 4). An example path, as found for *ϕ*_0_ = 0.1, is shown in Movie S1.

**FIG. 4.**
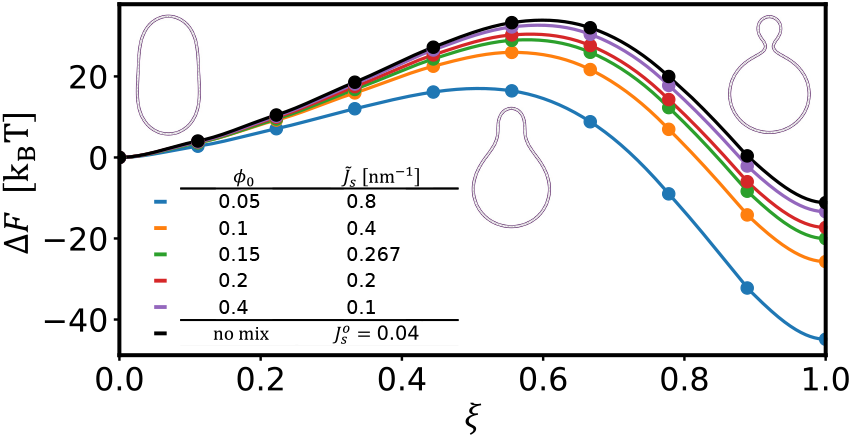
The budding free energy pathway along the NEB reaction coordinate for various vesicles with an average spontaneous curvature of 0.04 nm^*−*1^ in the outer leaflet, consistent with POPC vesicle parameters (see main text). Except for the non-mixing vesicle (black), the average spontaneous curvature arises from the additional lipid’s intrinsic curvature, 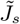, and its molar ratio, *ϕ*_0_. The vesicle transitions from a prolate shape (left) to a pear shape (middle) and ultimately forms a neck, separating into distinct MV and DV (right); these images represent the minimized path of the non-mixing system.

Overall, lipid mixing reduces the free energy difference both for the barrier and the final state. A slight increase in *f* ^mix^ via lipid demixing allows for a more substantial decrease in *f* ^el^ throughout budding (Fig. S8). The main energy components that change during this process are bending, which increases, and area stretching, which decreases (Fig. S9). The energy barrier decreases from 33.9 *k*_B_*T* for the non-mixed vesicle to 25.9 *k*_B_*T* for 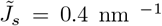 and *ϕ*_0_ = 0.1, which should increase the transition rate by several orders of magnitude. Notably, these energy barriers are close to those suggested for membrane fusion [24,26]. Therefore, when fusion occurs at the neck, the budding kinetics may determine its rate. This might be particularly relevant to biological processes, which typically involve asymmetric and heterogeneous membranes [4–6].

For non-ideal mixtures (Fig. S10), increasing the interaction parameter, *χ*, promoted budding to a certain extent. Specifically, increasing it from 0 to 2 further decreased the energy barrier for 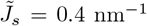 by 5.4 *k*_B_*T* to 20.5 *k*_B_*T*. Nevertheless, the process is extremely sensitive to the relative water content and the degree of area asymmetry (Figs. S11, S12), so even a small change in these factors may induce or inhibit it. This demonstrates the experimental limitations in terms of accuracy due to variations in vesicle properties within a sample or between fabrication methods.

Fig. 5A illustrates the compositional discrepancies between the DV and MV, Δ*ϕ* = ⟨*ϕ*⟩ _DV_ − ⟨*ϕ*⟩ _MV_, as well as the formation of domains, which arise due to increases in 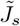 and *χ*; details in Section S7. The model parameters are as previously described, except for an initial relative water content of 90% and setting *ϕ*_0_ = 1*/*1.1 ≈ 0.0909 at the outer leaflet. Setting *ϕ*_0_ = 0.2 results in behavior similar to that detailed below (Fig. S7), provided that the average spontaneous curvature, 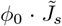, covers a similar range of values. When parameter values are low, no domain forms. However, if *χ* is sufficiently high, a circular domain forms at the top of the DV 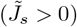 or a ring domain forms at the neck 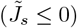. Further increases in the two parameters lead to the formation of a ring domain around the circular one, which eventually also becomes a ring domain. We expect a richer phase diagram for real vesicles, as the model only addresses cylindrically symmetric structures [37].

**FIG. 5.**
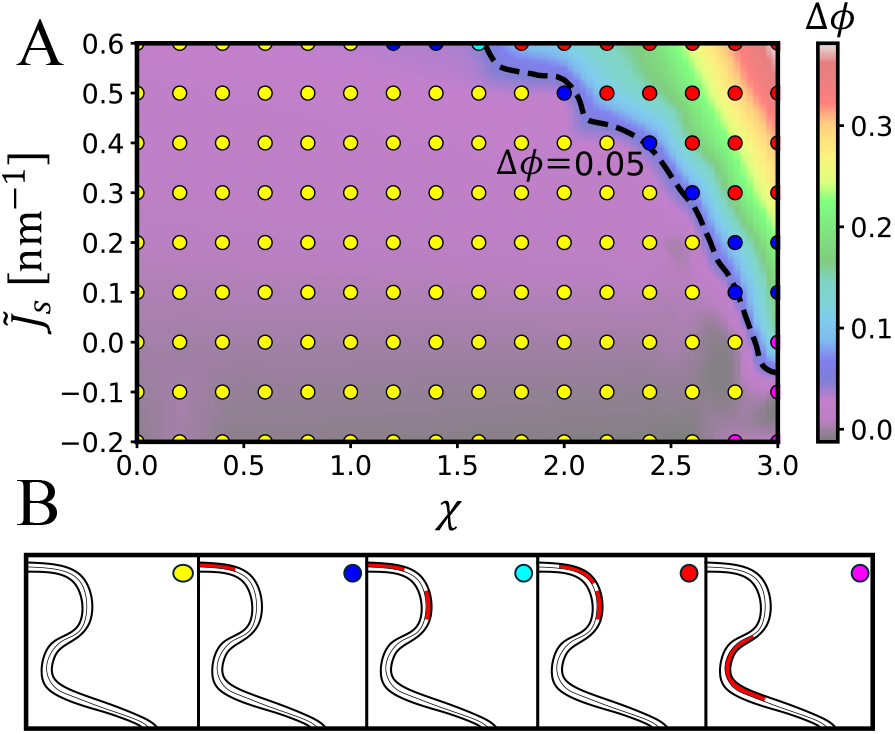
(A) The influence of the additional lipid’s spontaneous curvature, 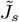, and the interaction parameter between the two lipids, *χ*, on compositional discrepancies between the DV and MV, Δ*ϕ*, and domain types in equilibrium structures. Some setups form no domain (B, yellow), while others form a single domain at the top of the DV (B, blue), sometimes with an additional ring domain (B, cyan). Other scenarios include two ring domains (B, red) or a single domain at the neck (B, magenta). Δ*ϕ* values were interpolated using a cubic spline and were less than 0.05 (dashed line) unless a domain formed.

With 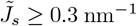, multiple ring domains appear. In this case, the additional lipid’s spontaneous curvature is larger than some micelle-forming surfactants [38]. Therefore, it is unlikely that these ring domains which contain more than 90% of a micelle-forming lipid will be stable in real vesicles but instead will separate from them as mixed micelles—a process which is not accounted for by the current model.

Generally, Δ*ϕ* is less than 0.05 (or 0.065 for *ϕ*_0_ = 0.2) unless a domain forms on the DV. Consequently, *f* ^mix^ and, by extension, the mixing entropy for sufficiently low *χ*, dominates over *f* ^el^ in its impact on the local composition. This agrees with other theoretical and experimental studies [30, 31, 39], where only a *relative* enrichment of less than 50% in some lipid component without phase separation in curved regions was observed. Notably, a recent experiment [17] under conditions similar to those modeled here reported up to a five-fold enrichment in lysolipid 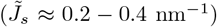 content in the DV. According to the model, such high enrichment should involve domain formation, which is unlikely in this case due to domain instability, as discussed above. This is also supported by MD simulations, which show that lysolipids in POPC should have *χ*≈ 0.6 ≪ 2 (Table S1), indicating a low tendency to phase separate.

In summary, the budding process in nanoscale vesicles was explored in a quantitative and controlled manner. The findings show that budding in such vesicles requires larger asymmetries. Additionally, careful modulation of membrane heterogeneity, area asymmetry, and osmotic conditions may allow for the regulation of budding dynamics. Finally, the composition of the DV is not substantially different from its MV unless phase separation occurs at the bud.

## I. ACKNOWLEDGMENTS

I thank D. Harries, P. Jungwirth, M. Javanainen, B. Fabian and M. I. Morandi for meaningful discussions and their insightful comments. I thank ChatGPT for aiding with refining the manuscript. I acknowledge the European Regional Development Fund OP RDE (project ChemBioDrug no. CZ.02.1.01/0.0/0.0/16 019/0000729) and the ERC Advanced Grant no. 101095957. by the European Research Council for support and computational resources.

