## Supplementary figures and images for "Lipid Demixing Reduces Energy Barriers for High Curvature Vesicle Budding"

### Movie S1

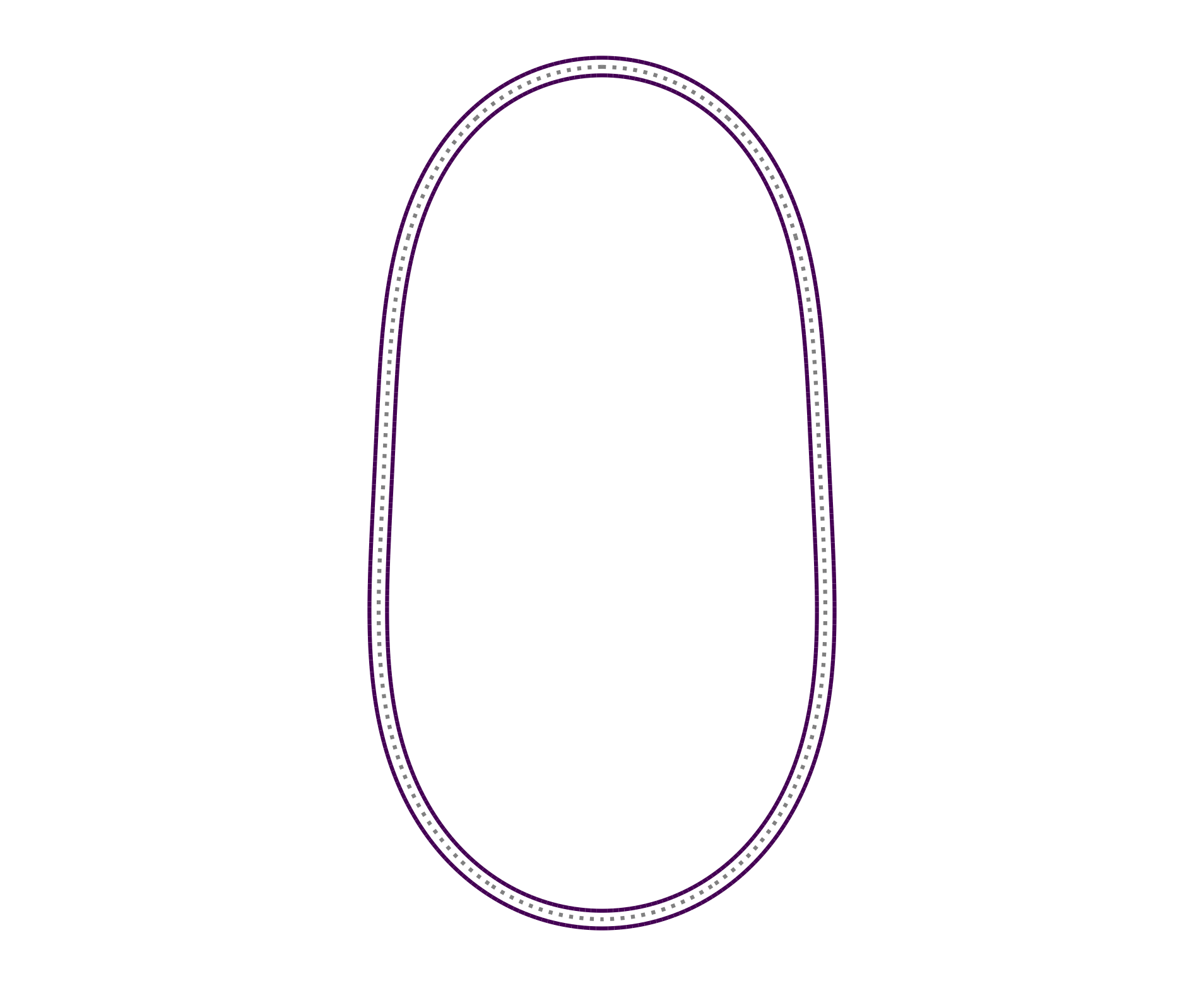
