## Supplementary material for "Lipid Demixing Reduces Energy Barriers for High Curvature Vesicle Budding": supplimentary information

Itay Schachter*

*Institute of Organic Chemistry and Biochemistry of the Czech Academy of Sciences, Flemingovo nám. 542/2, CZ-16000 Prague 6, Czech Republic and Institute of Chemistry, the Fritz Haber Research Center, and the Harvey M. Kruger Center for Nanoscience & Nanotechnology, The Hebrew University, Jerusalem 9190401, Israel*

(Dated: November 7, 2024)

### S1. THE ELASTIC MODEL IMPLEMENTATION

A finite-element method was employed to discretize the vesicle and approximate the free energy. Assuming the surfaces were cylindrically symmetric, they were represented in terms of $(r, z)$, with the default units being nanometers. The outer and inner leaflets and midplane surfaces, $S_o$, $S_i$, and $S_m$, were discretized into $N$ grid points: $\boldsymbol{x}^k_{o\backslash i\backslash m} = (r^k_{o\backslash i\backslash m}, z^k_{o\backslash i\backslash m})$ for $k = 1, \ldots, N$. The number $N$ was chosen to achieve a midplane resolution of $ds_m = 0.5$ nm for the results shown in Figures 5 and S7 due to the need of good description of the domain boundaries and $ds_m = 1$ nm for the rest. To maintain the resolution, frequent remeshing was performed using cubic spline interpolation of the grid, with a weak penalty imposed for changing the distances between successive midplane grid points. Detailed information can be found in the code itself (available here).

For $j = o, i$ the discretized lipid and tangent vector fields over $S_j$ are given by $\boldsymbol{D}^k_j = \boldsymbol{x}^k_m - \boldsymbol{x}^k_j = l^k_j \boldsymbol{n}^k_j$ and $\boldsymbol{\tau}^{k+\frac{1}{2}}_j = \boldsymbol{x}^{k+1}_j - \boldsymbol{x}^i_k$. Generally, half-indexed values are defined as the average of the adjacent full-index values, e.g., $\boldsymbol{n}^{k+\frac{1}{2}}_j = \frac{1}{2}(\boldsymbol{n}^{k+1}_j + \boldsymbol{n}^k_j)$, unless specified otherwise below. From these, the length $ds^{k+\frac{1}{2}}_j = |\boldsymbol{\tau}^{k+\frac{1}{2}}_j|$ and the area $dS^{k+\frac{1}{2}}_j = 2\pi r^{k+\frac{1}{2}}_j ds^{k+\frac{1}{2}}_j$ elements over $S_j$ are defined, along with the surface normal vector $\boldsymbol{N}^{k+\frac{1}{2}}$, which is the unit vector normal to the tangent vector and directed into the membrane.

The deformation modes are approximated as follows:

$$(\nabla \cdot \boldsymbol{n})^{k+\frac{1}{2}}_j = \frac{\boldsymbol{n}^{k+1}_j - \boldsymbol{n}^k_j}{ds^{k+\frac{1}{2}}_j} \cdot \boldsymbol{\tau}^{k+\frac{1}{2}}_j + \frac{\boldsymbol{n}^{k+\frac{1}{2}}_j}{r^{k+\frac{1}{2}}_j} \cdot \boldsymbol{e}_r,$$

$$\boldsymbol{t}^{k+\frac{1}{2}}_j = \frac{\boldsymbol{n}^{k+\frac{1}{2}}_j}{\boldsymbol{n}^{k+\frac{1}{2}}_j \cdot \boldsymbol{N}^{k+\frac{1}{2}}} - \boldsymbol{N}^{k+\frac{1}{2}},$$

$$\alpha^{k+\frac{1}{2}}_j = \frac{l_0 - l^{k+\frac{1}{2}}_j}{\sqrt{l_0 l^{k+\frac{1}{2}}_j}}$$

where $\boldsymbol{e}_r$ is the unit vector in the $r$-direction. The elastic free energy density is then estimated by:

$$f^{\mathrm{el},k+\frac{1}{2}}_j = K_C \left( (\nabla \cdot \boldsymbol{n})^{k+\frac{1}{2}}_j + J^{k+\frac{1}{2}}_{s,j} \right)^2 + \kappa_t \left| \boldsymbol{t}^{k+\frac{1}{2}}_j \right|^2 + K_A \left( \alpha^{k+\frac{1}{2}}_j \right)^2$$

where $J^{k+\frac{1}{2}}_{s,j}$ varies spatially in the case of binary mixtures, as detailed below.

*

For binary mixtures, an additional field, $\phi$, was required to represent the composition. Since $\phi$ must remain within the range $(0, 1)$, this can complicate numerical minimization methods. To address this, an auxiliary field, $\tilde{\phi}_j^k$, was defined over the interval $(-\infty, \infty)$ at the grid points of the mixed leaflet (hence, $\boldsymbol{x} = (r, z, \tilde{\phi})$). The actual value of $\phi_j^k$ was then computed as $\frac{1}{2}\left(\tanh(\tilde{\phi}_j^k) + 1\right)$. Also, $(\nabla\phi)_j^{k+\frac{1}{2}}$ was estimated as $\frac{\phi_j^{k+1} - \phi_j^k}{ds_j^{k+\frac{1}{2}}}$. Overall, this allowed the approximation of the mixing free energy density, $f_j^{\text{mix},k+\frac{1}{2}}$, as:

$$\frac{f_j^{\text{mix},k+\frac{1}{2}} \cdot (1 + \alpha_j^{k+\frac{1}{2}}) \cdot a_0}{k_{\text{B}}T} = \chi\phi_j^{k+\frac{1}{2}}\left(\phi_{j,0} - \phi_j^{k+\frac{1}{2}}\right) + \phi_j^{k+\frac{1}{2}} \log\left(\frac{\phi_j^{k+\frac{1}{2}}}{\phi_{j,0}}\right)$$

$$+ \left(1 - \phi_j^{k+\frac{1}{2}}\right) \log\left(\frac{1 - \phi_j^{k+\frac{1}{2}}}{1 - \phi_{j,0}}\right) + K\left|(\nabla\phi)_j^{k+\frac{1}{2}}\right|^2$$

To ensure the continuity of $\phi$, $K$ is set to 0.2. This choice results in a line tension of $0.25\,k_{\text{B}}T\,\text{nm}^{-1}$ for a linear increase in $\phi$ from 0 to 1 over 0.8 nm (approximately a lipid lateral length scale), which is close to the line tension observed in liquid-liquid phase coexistence [1]. This selection ensures that domains (i.e., regions with $\phi_0 \ll \phi < 1$) are of, at least, nanometric size while having a negligible effect on the energies and composition variances within a domain, as typically $|\nabla\phi|^2 \ll 0.01\,\text{nm}^{-2}$ within domains.

For binary mixtures, the local spontaneous curvature, $J_{s,j}^{i+\frac{1}{2}}$, was interpolated linearly between the spontaneous curvatures of the two components, $J_s^a$ and $J_s^b$, as:

$$J_{s,j}^{i+\frac{1}{2}} = (1 - \phi_j^{i+\frac{1}{2}})J_s^a + \phi_j^{i+\frac{1}{2}} J_s^b$$

As an extension of the original model by Ryham et al. [2–4], volume and composition constraints were incorporated. Specifically, the volumes of each leaflet, $V_o$ and $V_i$, and the water within the vesicle, $V_w$, were constrained to $V_o^0$, $V_i^0$, and $V_w^0$, respectively. These constraints were evaluated up to the first-order approximation in $ds_m$ and each was coupled to an appropriate Lagrange multiplier, $\lambda_m$ ($m = o, i, w$). Similarly, the composition of a leaflet, $\phi_j$ ($j = o, i$), was approximated and constrained to $\phi_{0,j}$ and coupled to a Lagrange multiplier, $\mu_j$, when necessary. Thus, the entire Lagrangian of the system is approximated as follows:

$$\mathcal{L}(\boldsymbol{X}) = \sum_{j=o,i} \sum_{k=1}^{N-1} \left( f_j^{\mathrm{el},k+\frac{1}{2}} + f_j^{\mathrm{mix},k+\frac{1}{2}} \right) ds_j^{k+\frac{1}{2}} + \sum_{j=o,i,w} \lambda_j \frac{V_j - V_j^0}{V_j^0} + \sum_{j=o,i} \mu_j (\phi_j - \phi_j^0)$$

where $\boldsymbol{X}$ is the hyper-vector $(\boldsymbol{x}_j^i)_{j=u,l,m}^{i=1,\ldots,N}$.

The structural optimization provided the equilibrium structures, achieved when $\nabla_{\boldsymbol{X}} \mathcal{L} = 0$. As $\boldsymbol{X}$ represents a saddle point of $\mathcal{L}$, reaching this point via standard functional minimization schemes is challenging. To address this, a modified version of the augmented Lagrangian method [5] was applied. In this method, the term $\sum_{j=o,i,w} c_j \left( \frac{V_j - V_j^0}{V_j^0} \right)^2$ was added to the Lagrangian, with the quadratic penalty coefficient $c_j$ chosen large enough to transform the saddle point into a minimum in the modified Lagrangian. Specifically, $\lambda_j$ and $c_j$ were updated after each optimization step. Initially, $c_j$ was set to $5 \cdot 10^4$, and then multiplied by 0.999 after each optimization step until reaching $10^4$. For the composition constraints, the initial quadratic penalty coefficients were set to $5 \cdot 10^6$, and the corresponding Lagrange multiplier was denoted by $\mu_k$, with $k$ potentially being set to 1 or 2.

Each optimization step consisted of 100 sub-steps, using the `FIRE2.0` algorithm [6, 7] with parameters $\alpha_{in} = 0.25$, $dt_{in} = t_{min} = 10^{-5}$, $t_{\mathrm{max}} = 10^{-1}$, $N_{\mathrm{del}} = 5$, $\alpha_{dec} = 0.99$, $dt_{inc} = 1.1$, $dt_{dec} = 0.5$, and $N_{\mathrm{ple,max}} = \infty$ (i.e., not implemented). The gradient $\nabla_{\boldsymbol{X}} \mathcal{L}$ was evaluated numerically via a centered finite difference scheme with $\Delta x = 10^{-5}$. Optimization halted when the energy difference between subsequent steps was lower than $2.5 \cdot 10^{-4}\,\mathrm{pN} \cdot \mathrm{nm} \approx 6 \cdot 10^{-5}\,\mathrm{k_B T}$ and constraints were satisfied up to an error of $10^{-5}$. An example of an equilibrated configuration is shown in Figure S1

The model was implemented in Julia [8] and is available here.

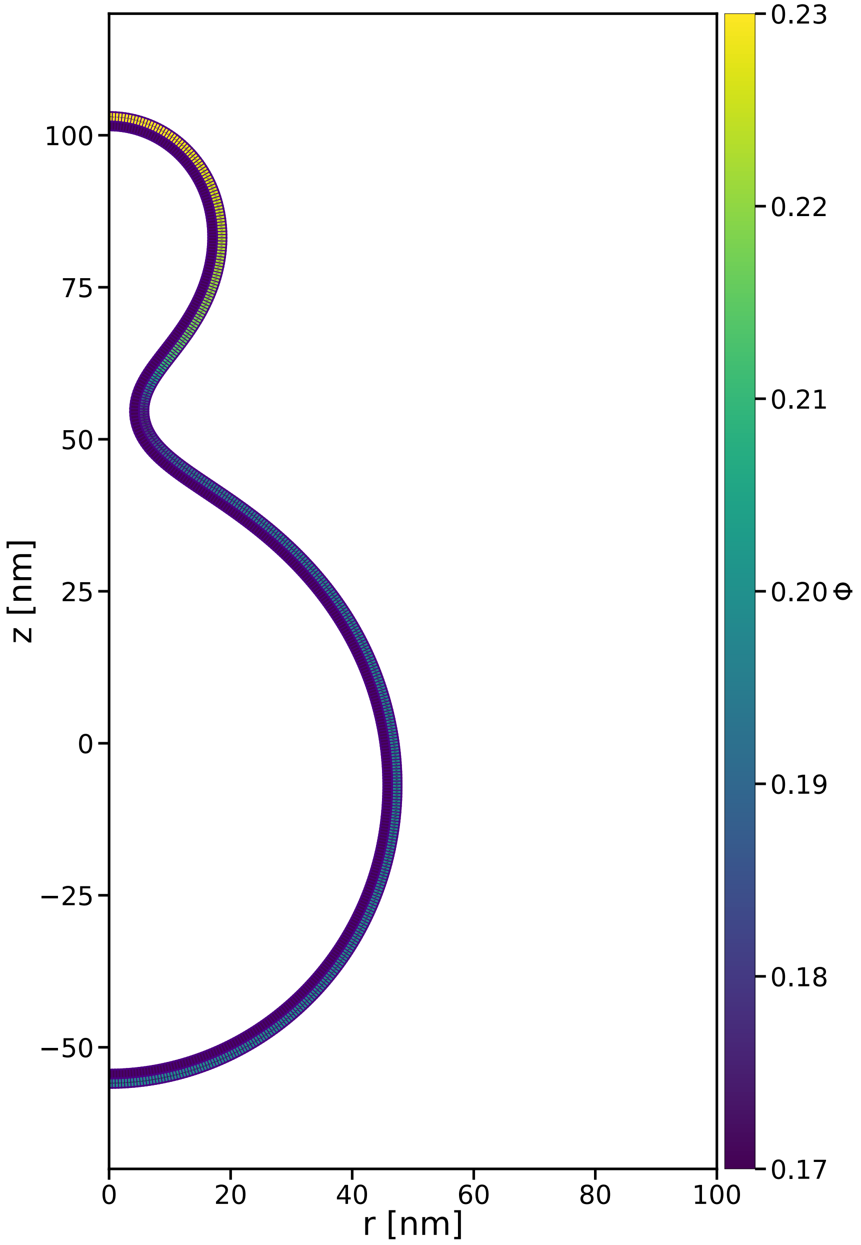


FIG. S1. Example for an optimized configuration of a budded vesicle with a binary mixture.

### A. Model and Implementation Validation

Due to the need of validation of the implementation and concern which stemmed from discrepancies between the results of the current model and those reported for the Area Difference Elasticity (ADE) model [9] (see Section S2 and the main text). Specifically, the current model behaved similarly to the ADE model with a one-fourth of the value of $\alpha = \frac{K_a l_0^2}{\pi K_C}$. Thus, I mainly verified that: 1) $K_a$ and $K_C$ are well-defined as well as the volume constraints. 2) The current implementation recovers some previously published results [3]. To this aim, the following three test cases were applied.

#### 1. *Single Monolayer Stretching*

To validate that area stretching behaves as it should, I made a discoidal monolayer of 50 nm radius ($= r$) with $K_a = 1\ \frac{\mathrm{pN}}{\mathrm{nm}}$, with different volumes, such that $\alpha = (a - a_0)/\sqrt{a_0 a}$ obtained the values $\alpha = 0, 0.01, 0.1, 1.5$, the resulting energies were $0, 0.3886, 35.69, 654.30$ pN nm. All these values are same as the one obtain by the analytical expression, $0.5\pi r^2 K_a \alpha^2$ up to 0.1% error.

#### 2. *Bending to a Half Sphere*

To validate that bending energy and volume constraints behave as they should, I made an initially flat discoidal bilayer with a free edge, defining $R = 20$ nm, the volume of the outer leaflet was set to $V_o = 0.5h_0\pi(R + 0.5h_0)^2$ and its spontaneous curvature was set to $J_s^o = \frac{2}{R+h_0}$. Similarly, the inner leaflet had volume of $V_i = 0.5h_0\pi(R - 0.5h_0)^2$ and $J_s^i = -\frac{2}{R-h_0}$. Under these conditions, the bilayer should bend into a half sphere, without any area stretch or tilt deformations, with and a total energy of $-0.5K_C(J_s^o A_o + J_s^i A_i)$, given that $A_o = 2\pi(R+h_0)^2$ and $A_i = 2\pi(R-h_0)^2$ the energy of such a half sphere is $8\pi K_C$ which for $K_C = 10\ k_\mathrm{B}T$ equals to $-251.32\ k_\mathrm{B}T$. As expected, the membrane bent into a half sphere with an energy value $-251.56\ k_\mathrm{B}T$ which is close to the analytical value.

#### 3. Toroidal Membrane Pore

The implementation was validated by recovering the energies for a toroidal membrane pore, as reported in Figures 4B and 7A in Ref. [3], after accounting for the authors' choice of orientation.

### S2. MISCALCULATION IN THE ORIGINAL ADE MODEL PAPER

Following the discussion in the main text, as $\frac{l_0}{R_a} \to 0$, we expect the current model to recover the coexistence curve in the $(\nu, m_0)$ plane as predicted by the ADE model, given a specific value of $\alpha = \frac{K_a l_0^2}{K_C \pi}$. However, when setting $l_0 = 0.5$ nm and $\alpha = 4$ (Fig. 3) or $\alpha = 16$ (Fig. S2), the resulting curves are notably closer to those found in the ADE model for $\alpha = 1$ or $\alpha = 4$, respectively.

Since the current model implementation behaves as expected (Section S1 A), this suggests that the $\alpha$ incorporated into the numerical calculations in Ref. [9] was multiplied by a factor of $\frac{1}{4}$.

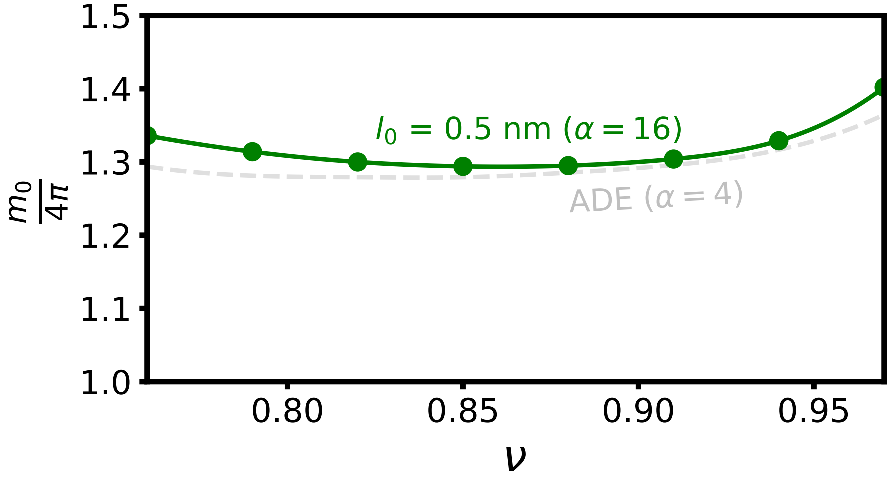


FIG. S2. Phase diagram of the elastic model in the $(\nu, m_0)$ plane involving budding with the coexistence curve being compared between the ADE model for *alpha* = 4 as found in Ref. [9] (grey dashed line) and the current model (green line) for a 50 nm vesicle with the following elastic parameters: $l_0 = 0.5$ nm, $K_C = 10$ $\mathrm{k_B T}$, $\kappa_t = 10 k_B T$ nm$^{-2}$ and $K_a = \frac{\alpha \pi K_C}{l_0^2}$ with $\alpha = 16$. Above the coexistence curve the vesicle is budded and below it has a prolated shape.

### S3. CONSTRUCTION OF THE $\frac{m_0}{4\pi}$-$\nu$ PHASE DIAGRAM

The coexistence curves shown in the phase diagrams in Figs 3 and S2 were evaluated via a cubic spline, connecting values calculated at $\nu = [0.76, 0.79, 0.82, 0.85, 0.88, 0.91, 0.94, 0.97]$. The curve was found in each $\nu$ by optimizing the structure of initially budded and initially prolate vesicles for $\frac{m_0}{4\pi}$ of different values in jumps of 0.1. For each $m_0$, assuming that both structures maintained their phase, $\Delta E$ was evaluated. From the three closest points to $\Delta E = 0$, a quadratic interpolation was made to find the value of $m_0$ which gives $\Delta E = 0$, i.e. a point on the coexistence curve, as shown in Fig. S3. In cases where the coexistence curve required extrapolation, smaller jumps in $m_0$ were made to ensure the usage of interpolation.

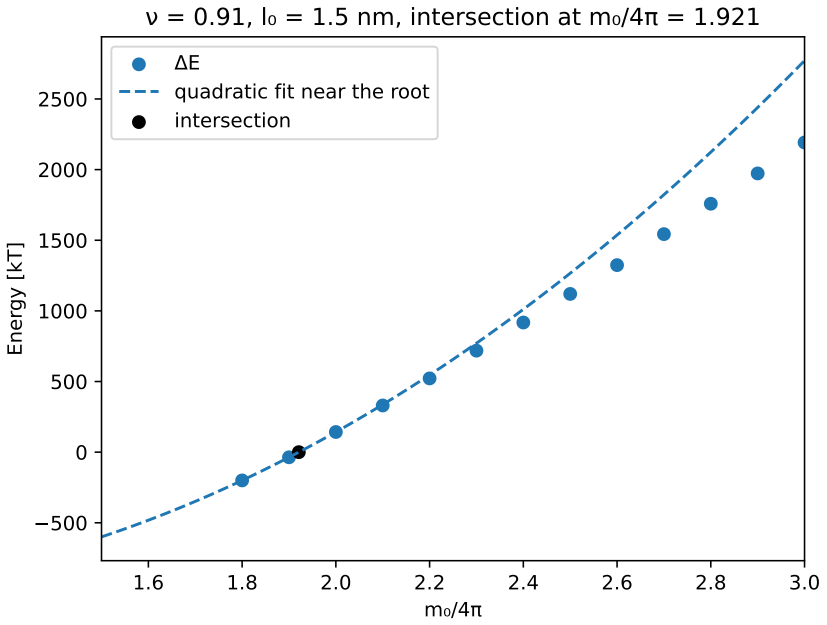


FIG. S3. Example of the interpolation scheme for finding the $m_0$ on the coexistence curve for $\nu = 0.91$. Further details in the text.

### S4. JUSTIFICATION FOR THE COMPOSITIONAL FREE ENERGY FUNCTIONAL FOR TWO COMPONENT MEMBRANES

The suitability of employing the free energy functional from Eq. 2 is demonstrated in this section. We begin by deriving a relationship between the local composition variance and the applied external potential for the selected $f^{\mathrm{mix}}(\phi)$. Subsequently, Molecular Dynamics

(MD) simulations were conducted with this external potential. The interaction coefficient $\chi$ was determined by fitting the composition profiles, yielding an excellent goodness of fit and indicating that the approximation is valid across a wide range of concentrations for various binary mixtures.

#### A. Compositional Variations under External Potential

Consider a flat, finite rectangular membrane defined for $\boldsymbol{x} = (x, y) \in [0, L_1] \times [0, L_2]$. Assume that the free energy density $f$ is a functional of $\phi$. Under the influence of an axisymmetrical external potential $\phi(\boldsymbol{x}) \cdot u(x)$, the total energy function is

$$F[\phi] = L_2 \int_0^{L_1} \left( f(\phi(x)) + \phi(x) \cdot u(x) \right) dx,$$

neglecting the dependence on $y$ due to the symmetry of $u$. Simulations of lipid membranes are typically performed with a fixed number of lipids, ensuring that the average component ratio

$$E[\phi] = \frac{1}{L_1} \int_0^{L_1} \phi(x) dx$$

remains constant. This constraint is enforced using an appropriate Lagrange multiplier, $\lambda$, and the condition $G[\phi] = E[\phi] - \phi_0 = 0$ for some $\phi_0$. Therefore, the functional

$$\bar{F}[\phi] = F[\phi] + \lambda G[\phi]$$

is minimized, leading to:

$$\frac{\partial \bar{F}}{\partial \phi} = \frac{\partial f}{\partial \phi} + u(x) + \lambda = 0.$$

By inserting $f^{\text{mix}}$, we obtain:

$$u(x) \cdot a_0 = k_{\text{B}} T [2\chi\phi(x) - \ln\left(\frac{\phi(x)}{1 - \phi(x)}\right) + \chi - \lambda] \quad \text{(S1)}$$

When $u(x_0) = 0$ for a certain $x_0$, denoted by $\phi_a = \phi(x_0)$, it follows that

$$\lambda = k_{\text{B}} T [2\chi\phi_a - \ln\left(\frac{\phi_a}{1 - \phi_a}\right) + \chi],$$

leading to:

$$u(x) \cdot a_0 = k_{\text{B}} T [2\chi(\phi(x) - \phi_a) - \ln\left(\frac{\phi(x)}{1 - \phi(x)}\right) + \ln\left(\frac{\phi_a}{1 - \phi_a}\right)] \quad \text{(S2)}$$

#### B. Comparison with MD Simulations

To assess the validity of the $f^{\mathrm{mix}}(\phi)$, molecular dynamics simulations were conducted using the Dry Martini force field [10]. Initial configurations were prepared as follows: a box of 100 nm x 20 nm x 20 nm with a POPC bilayer on the XY plane was created using INSANE [11]. Subsequently, an in-house script utilizing MDAnalysis [12] was used to replace 40% percentage of the POPC lipids in each leaflet with the lipids described in Table S1, giving a partial molar ratio $\phi_0 = 0.4$. Using Gromacs 5.1.5 [13], each simulation ran for a total of 24 microseconds, using the first 16 microseconds as equilibration. The simulation setup mirrored that of Ref.[10] at T=298 K, except for simulations including DPPC, which were conducted at T=320 K. Additionally, a simulation with a pure POPC bilayer was performed under the same conditions for 180 ns after an equilibration period of 20 ns. The area per lipid ($a_0$) for the Dry Martini POPC was evaluated as APL$= 0.6310 \pm 0.0001$ nm$^2$.

An inverted flat-bottom potential with the distance function $r(x) = |x - x_0|$, where $x_0$ is set to the center of the simulation box, was applied to the phosphate group of the secondary lipids; as well as an weak flat-bottom potential in the $z$-direction to ensure membrane stability. The potential function is given by $u(x) = -K\frac{r^2}{r_{\mathrm{max}}^2}$ with $K = 3\ k_{\mathrm{B}}T$ and $r_{\mathrm{max}} =$ 22.5 nm. Composition profiles were sampled over the last 8 microseconds, binned into approximately 1 nm width bins, and $\phi_a$ was averaged for $|x| > 30$ nm. Given $u(x)$, $a_0$, and $\phi(x)$, Eq.S2 was fitted over $|x| \in [4, 18]$ nm, resulting in the interaction parameter $\chi$ with the $R^2$ of the fit. The goodness of fit for POPC:DOPC is shown in Figure S4, and the high $R^2$ values indicate that the $f^{\mathrm{mix}}$ is a reliable approximation for a range of binary lipid mixtures. While setting $\phi_0$ to a lower value might provide a range for $\phi(x)$ that is more consistent with its values that were chosen in some parts of this paper, it proved problematic because the results are highly sensitive to $\lambda$. This sensitivity is especially pronounced when $\phi_a$ is close to zero, as errors in $\phi_a$ significantly impact $\lambda$, which approaches $-\infty$ at $\phi_a = 0$.

TABLE S1. MD simulations details and analysis results

| Membrane Composition (Ratio) | T [K] | $\phi_{\min}$[14] | $\phi_{\max}$[14] | $\chi$ $[\frac{k_B T}{nm^2}]$ | $R^2$ |
|---|---|---|---|---|---|
| POPC:DOPC (6:4) | 298 | 0.32 | 0.71 | $0.74 \pm 0.01$ | 0.998 |
| POPC:DPPC (6:4) | 320 | 0.30 | 0.79 | $1.01 \pm 0.05$ | 0.971 |
| POPC:POPE (6:4) | 298 | 0.30 | 0.73 | $0.68 \pm 0.04$ | 0.995 |
| POPC:LPC (6:4) | 298 | 0.30 | 0.66 | $0.61 \pm 0.01$ | 0.998 |

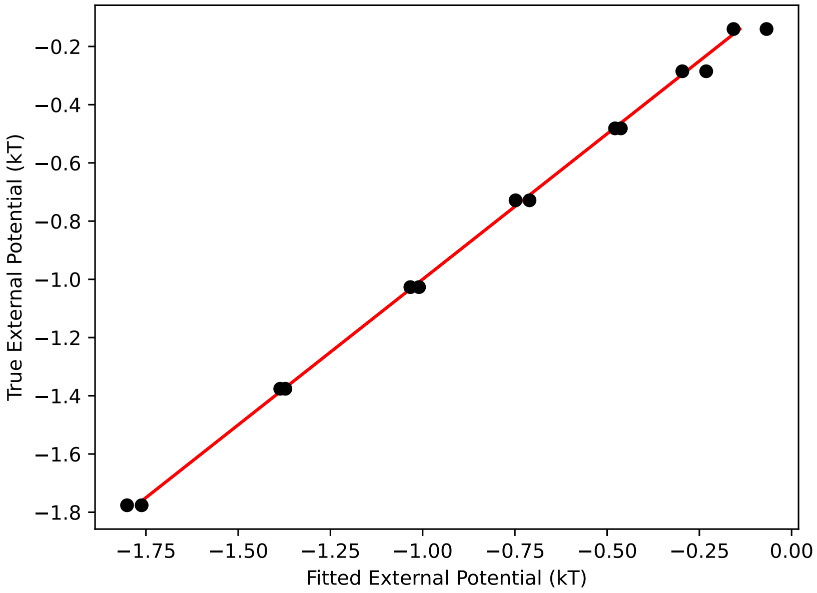


FIG. S4. A goodness-of-fit plot compares the fitted external potential with the true one for the POPC:DOPC membrane over the fitted region (black circles). Ideally, this plot should align with the line $y = x$ (red line).

### S5. MINIMAL FREE ENERGY PATH USING A MODIFIED NUDGED ELASTIC BAND METHOD

A minimal (free) energy path minimizes the height of the budding transition energy barrier, connecting the "opened" state to the "budded" state. Moving down (up) a least energy path corresponds to the steepest gradient descent (ascent); it ignores any energy dissipation and does not account for fluctuations.

Numerically, the free energy path is discretized into $M$ connected images; the realization of this connectivity is method-dependent [15, 16]. Previously [4], a simplified string method [16] was employed to probe the energy path of membrane fusion. This method involves frequent interpolation between all images for each grid point, preferably at every optimization step. In the current model, each image $m$ has a different number of grid points, $N_m$, to preserve resolution. Consequently, each image must be remeshed before interpolation, which hinders structural relaxation. To reduce the frequent remeshing, only two representative $(r, z)$ grid points in the outer leaflet, represented by the hyper-vector $\boldsymbol{X}_m$ for each image $m$, were connected between images. Specifically, these points were: (1) the point at an arc-length distance $d$ from the head of the bud, chosen to be at the middle of the neck region of the last image, and (2) a point in the middle at a distance of $\frac{d}{2}$ from the head of the bud, roughly located at the region with a maximal $r$-value in the bud. Furthermore, to avoid arbitrary translations of images, a point at an arc-length distance of 20 nm from the bottom of the vesicles was anchored to $z = 0$. This approach allowed for setting a continuous transition path without frequent remeshing. However, the simplified string method was still unstable under this scheme and tended to crash.

To enhance stability, a modified version of the nudged elastic band (mNEB) method [15, 17, 18] was applied. First, the tangent direction to the path at the $m^{\text{th}}$ image is defined as an energy-dependent combination of the forward and backward tangents, $\boldsymbol{\tau}_m^+ = \boldsymbol{X}_{m+1} - \boldsymbol{X}_m$ and $\boldsymbol{\tau}_m^- = \boldsymbol{X}_m - \boldsymbol{X}_{m-1}$ [17]:

$$
\boldsymbol{\tau}_m = \begin{cases} \boldsymbol{\tau}^{+} & \text{if } \mathcal{L}_{m+1} > \mathcal{L}_m > \mathcal{L}_{m-1} \\ \boldsymbol{\tau}^{-} & \text{if } \mathcal{L}_{m+1} < \mathcal{L}_m < \mathcal{L}_{m-1} \\ \boldsymbol{\tau}^{+}\Delta\mathcal{L}_m^{\max} + \boldsymbol{\tau}^{-}\Delta\mathcal{L}_m^{\min} & \text{otherwise and } \mathcal{L}_{m+1} > \mathcal{L}_{m-1} \\ \boldsymbol{\tau}^{+}\Delta\mathcal{L}_m^{\min} + \boldsymbol{\tau}^{-}\Delta\mathcal{L}_m^{\max} & \text{otherwise} \end{cases}
$$

where $\mathcal{L}_m^{\max} = \max(|\mathcal{L}_{m+1} - \mathcal{L}_m|, |\mathcal{L}_m - \mathcal{L}_{m-1}|)$ and $\mathcal{L}_m^{\min} = \min(|\mathcal{L}_{m+1} - \mathcal{L}_m|, |\mathcal{L}_m - \mathcal{L}_{m-1}|)$ with $\mathcal{L}_j$ being the current Lagrangian value at the $j^{\text{th}}$ image. The method only modifies the force at the connected grid points for each image $m$ (except for the initial and final images, $m = 1, M$):

$$
\boldsymbol{F}_{\boldsymbol{X}_m} = -\nabla_{\boldsymbol{X}_m}\tilde{\mathcal{L}}_m = -\left(\nabla_{\boldsymbol{X}_m}\mathcal{L}_m\right)_{\perp} + \left(\boldsymbol{F}_m\right)_{\parallel} + f(\theta_m)\left(\boldsymbol{F}_m\right)_{\perp}
$$

The first term is the normal component to the energy path of the original gradient of the Lagrangian of the image. The second term is a spring penalty term, tangent to the energy path, aimed at keeping the distance between subsequent $\boldsymbol{X}_j$s equal, defined as $\left(\boldsymbol{F}_m\right)_{\parallel} = k\left(|\boldsymbol{\tau}_m^{+}| - |\boldsymbol{\tau}_m^{-}|\right)\hat{\boldsymbol{\tau}}_m$ with a spring constant $k$ of 50 $\frac{\text{pN}}{\text{nm}}$. The spring constant does not affect the resultant energy path [15] and its value was chosen to maintain stability. The last term, which improves the stability of mNEB at curved regions of the energy path, is the normal component of the spring penalty, $\boldsymbol{F}_m = k\left(\boldsymbol{\tau}_m^{+} - \boldsymbol{\tau}_m^{-}\right)$, which is activated by a switching function $f(\phi_m) = \frac{1}{2}\left(1 + \cos\left(\pi\cos(\phi_m)\right)\right)$ where $\cos(\phi_m) = \frac{\boldsymbol{\tau}_m^{+}\cdot\boldsymbol{\tau}_m^{-}}{|\boldsymbol{\tau}_m^{+}||\boldsymbol{\tau}_m^{-}|}$. The constraints were updated separately for each image, as described in Ref. [19]. Finally, a smooth interpolat 2) without the mixing entropy ion of the energy path is obtained from the tangential forces at each image[17].

The structure optimization was performed similarly to the method described in Section S1 for individual configurations. Before optimizing the energy path, the initial and final states were thoroughly minimized. The number of images was set to $M = 10$, as increasing the number of images did not significantly impact the resulting energy barrier (less than 0.2 $k_B T$), as shown in Figure S5. The optimization process was halted when the energy change of any image between subsequent steps was less than $0.5 \cdot 10^{-4}$ pN nm and all the constraints were held up to $10^{-4}$.

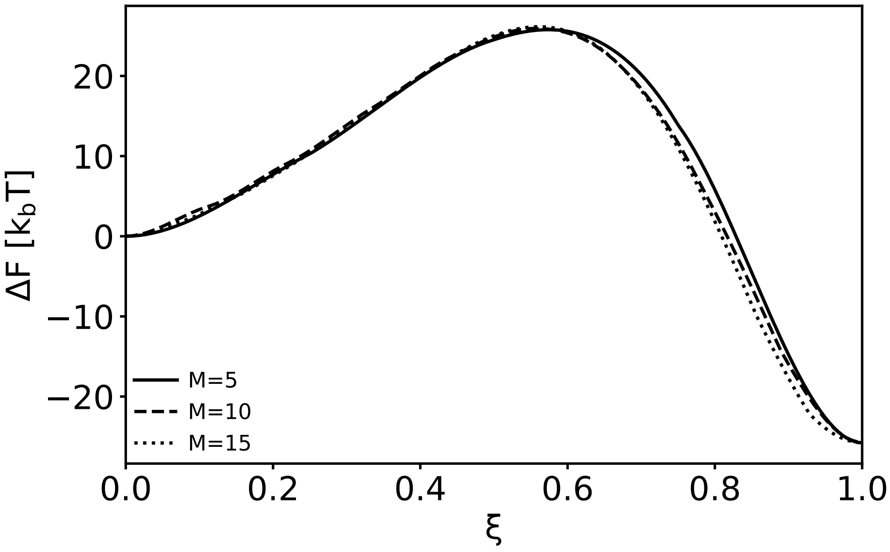


FIG. S5. The budding free energy path over the NEB reaction coordinate for a vesicle, resulting from the usage of different number of images. Parameters match a POPC vesicle (see main text) except the outer leaflet being a binary mixture with the second lipid's intrinsic curvature, $\tilde{J}_s = 0.4$ nm$^{-1}$, of a molar ratio, $\phi_0 = 0.1$. The path starts from a prolated shape vesicle ($\xi = 0$) and ends with a budded vesicle ($\xi = 1$)

### S6. CONSTRUCTION OF THE $\tilde{J}_s$–$\chi$ PHASE DIAGRAM

To efficiently evaluate the phase diagram presented in Figure 5 of the main text within reasonable computational time, required choosing initial structures close to the equilibrium ones. Grid points were sampled at $(J_s, \chi) \in (-0.2, -0.1, 0, ..., 0.6) \times (0, 0.2, 0.4, ..., 3)$, with $J_s$ in units of nm$^{-1}$. The configurations were optimized for each pair of $(J_s, \chi)$ using previously equilibrated structures with similar parameter values. Different trajectories were employed to achieve the true minimized equilibrium structure out of several locally minimal structures (refer to Figure S6 for detailed sampling trajectories):

- Path A started from an optimized structure at $(0, 0)$ without any domain.
- Path B began with a configuration featuring 2 ring domains on the bud at $(0.6, 3)$.
- Path C initiated from a circular domain on the top of the bud at $(0.1, 3)$.
- Path D started with a single ring domain at the neck region at $(-0.1, 3)$.

For $J_s = 0.6$ and $\chi = 1.2, 1.4, 1.6$, a finer mesh resolution of 0.3 nm was used because different locally minimized structure had similar energies (up to 1.5 $k_B T$) and the presence of smaller domains required a more precise characterization.

The equilibrium structure for each $(J_s, \chi)$ point was determined as the configuration with the lowest energy among those optimized for it.

A similar analysis was conducted for $(J_s, \chi) \in (-0.05, 0, ..., 0.3) \times (0, 0.2, 0.4, ..., 3)$ at $\phi_0 = 0.2$ (see Figure S7). This narrower phase space was selected to ensure that the average spontaneous curvature, $\phi_0 \cdot \tilde{J}_s$, remained of the same order of magnitude; for larger values, the overall stability of the bilayer phase is questionable.

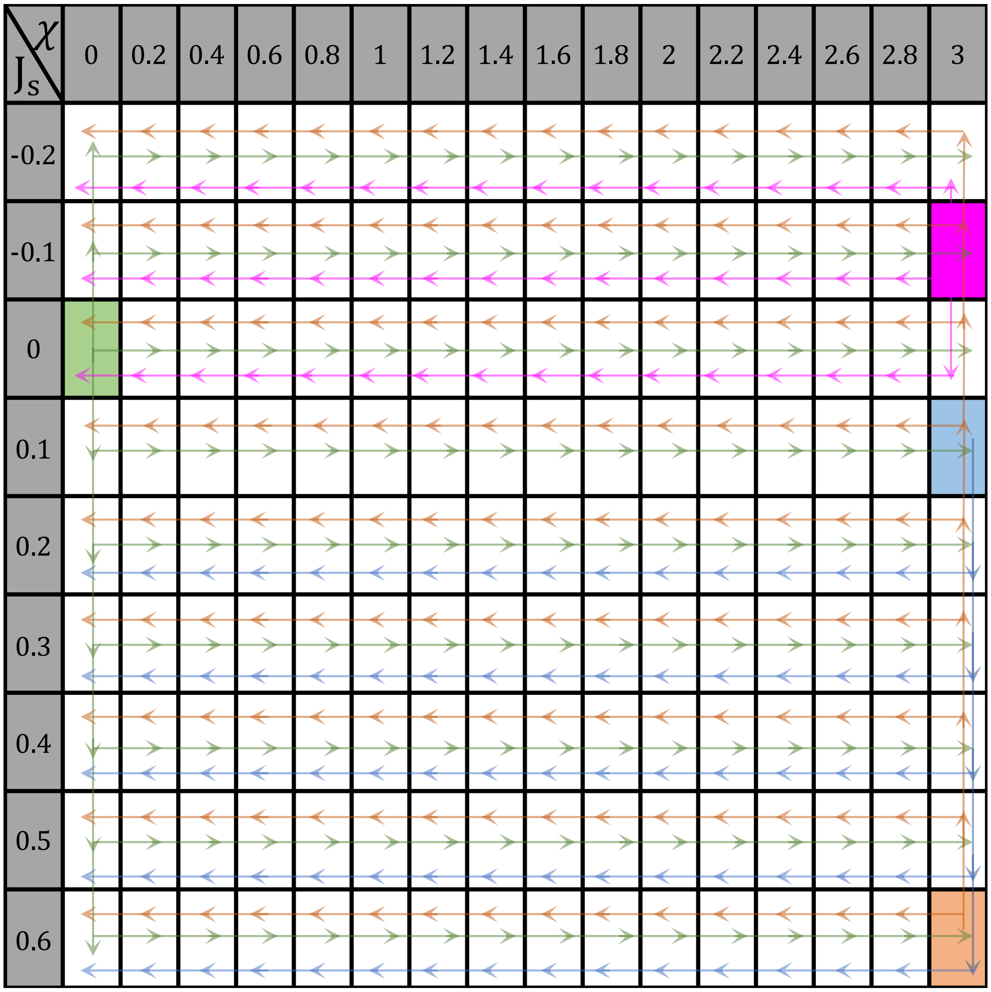


FIG. S6. Schematic representation of the sampling trajectories across the parameter space ($\chi$ vs $\tilde{J}_s$) for constructing the phase diagram, as discussed in the text. Various paths (A-green, B-orange, C-blue, D-purple) were taken over the phase diagram grid.

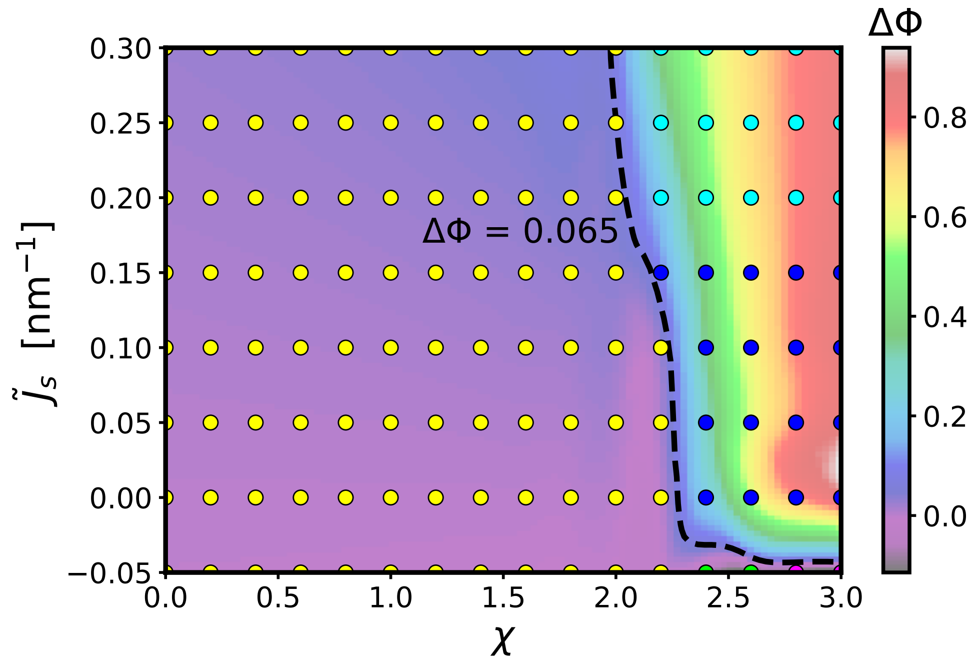


FIG. S7. Influence of the spontaneous curvature of the additional lipid, $\tilde{J}_s$, and the interaction parameter between the two lipids, $\chi$, on the compositional discrepancies between the DV and MV, $\Delta\phi$, and the types of domains formed in the equilibrium structures of the sampled parameters (points) in the setup detailed in the main text. In some cases, no domain forms (yellow), while in others a single domain forms at the top of the DV (blue), with a possible additional ring domain forming around it (cyan). In other scenarios, a single domain formed at the neck (magenta) or the MV (green). The values of $\Delta\phi$ were interpolated using a cubic spline and were less than 0.065 (dashed line) unless a domain formed.

### S7. SUPPLEMENTARY FIGURES

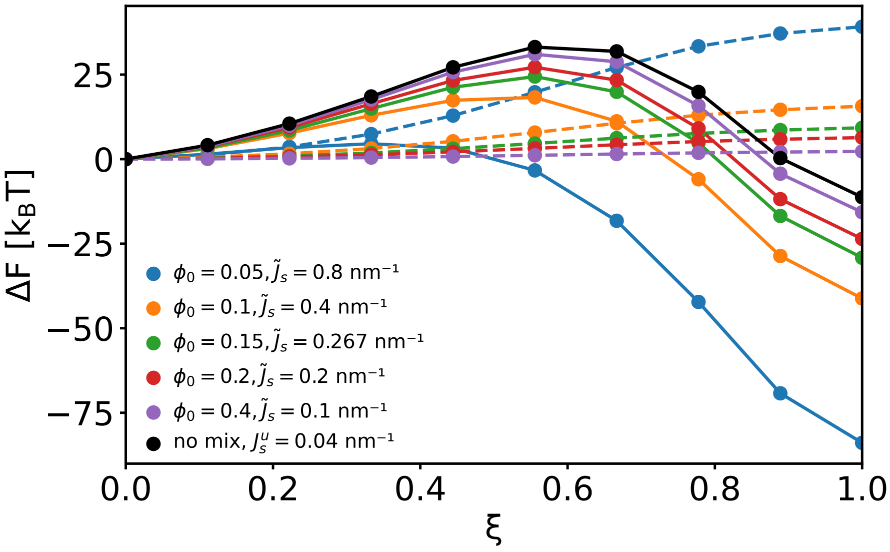


FIG. S8. The elastic (solid lines) and mixing (dashed lines) free energies throughout the budding path for the different vesicles as described in the main text and Fig. 4.

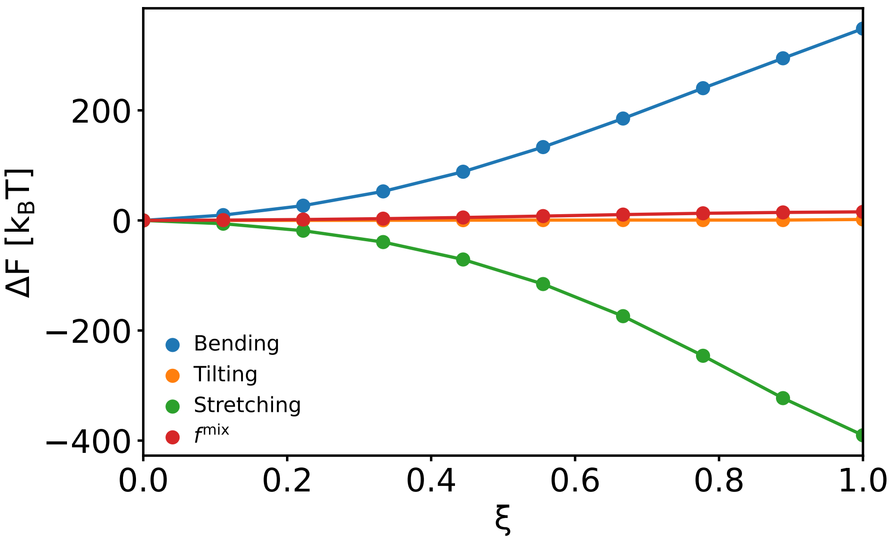


FIG. S9. The different energetic contributions throughout the budding path for the vesicle with $\phi_0 = 0.1$ and $\tilde{J}_s = 0.4$ nm$^{-1}$ as described in the main text.

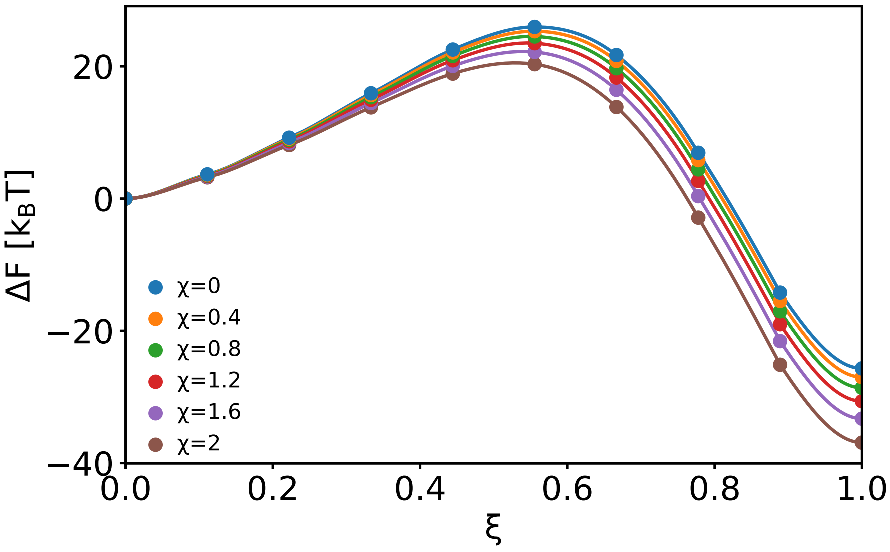


FIG. S10. The effect of non-ideal mixing, represented by the $\chi$ parameter, on the budding energy path for the vesicle with $\phi_0 = 0.1$ and $\tilde{J}_s = 0.4$ nm$^{-1}$ as described in the main text.

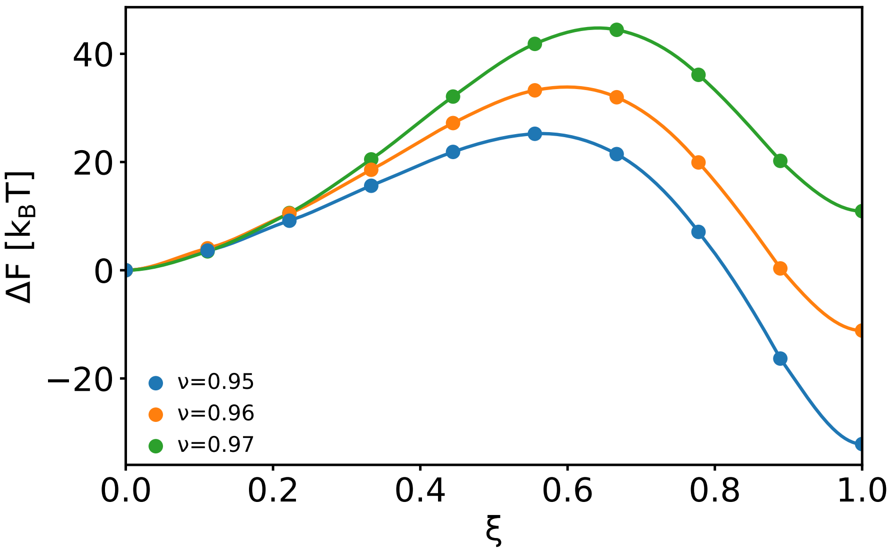


FIG. S11. The effect of the initial vesicle relative hydration, represented as $\nu$, on the budding energy path for the vesicle with $\phi_0 = 0.1$ and $\tilde{J}_s = 0.4$ nm$^{-1}$ as described in the main text.

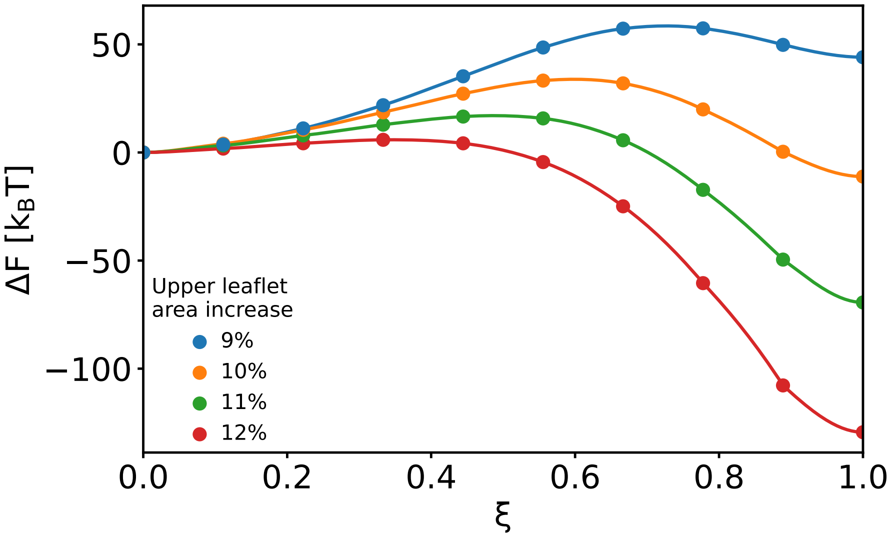


FIG. S12. The effect of the outer leaflet area increase from its initial, relaxed, value on the budding energy path for the vesicle with $\phi_0 = 0.1$ and $\tilde{J}_s = 0.4$ nm$^{-1}$ as described in the main text.

[13] M. J. Abraham, T. Murtola, R. Schulz, S. Páll, J. C. Smith, B. Hess, and E. Lindahl, Gro-

macs: High performance molecular simulations through multi-level parallelism from laptops to supercomputers, SoftwareX **1**, 19 (2015).

[14] In the fitting region.
